# A transcriptomic atlas of the mouse cerebellum reveals regional specializations and novel cell types

**DOI:** 10.1101/2020.03.04.976407

**Authors:** Velina Kozareva, Caroline Martin, Tomas Osorno, Stephanie Rudolph, Chong Guo, Charles Vanderburg, Naeem Nadaf, Aviv Regev, Wade Regehr, Evan Macosko

## Abstract

The cerebellum is a well-studied brain structure with diverse roles in motor learning, coordination, cognition, and autonomic regulation. Nonetheless, a complete inventory of cerebellar cell types is presently lacking. We used high-throughput transcriptional profiling to molecularly define cell types across individual lobules of the adult mouse cerebellum. Purkinje and granule neurons showed considerable regional specialization, with the greatest diversity occurring in the posterior lobules. For multiple types of cerebellar interneurons, the molecular variation within each type was more continuous, rather than discrete. For the unipolar brush cells (UBCs)—an interneuron population previously subdivided into two discrete populations—the continuous variation in gene expression was associated with a graded continuum of electrophysiological properties. Most surprisingly, we found that molecular layer interneurons (MLIs) were composed of two molecularly and functionally distinct types. Both show a continuum of morphological variation through the thickness of the molecular layer, but electrophysiological recordings revealed marked differences between the two types in spontaneous firing, excitability, and electrical coupling. Together, these findings provide the first comprehensive cellular atlas of the cerebellar cortex, and outline a methodological and conceptual framework for the integration of molecular, morphological, and physiological ontologies for defining brain cell types.

---

The cerebellar cortex is composed of the same basic circuit replicated thousands of times. Mossy fibers from many brain regions excite granule cells (GCs) that in turn excite Purkinje cells (PCs), the sole outputs of the cerebellar cortex. Powerful climbing fiber synapses originating in the inferior olive excite PCs and regulate synaptic plasticity. Additional circuit elements include inhibitory interneurons such as MLIs, purkinje layer interneurons (PLIs), Golgi cells (GoCs), and excitatory UBCs. There is a growing recognition that cerebellar circuits exhibit regional specializations, such as a higher density of UBCs or more prevalent PC feedback to GCs in some lobules. Molecular variation across regions has also been identified, such as the parasagittal banding pattern of alternating PCs with high and low levels of *Aldoc* expression^1^. However, the extent to which cells are molecularly specialized in different regions is poorly understood.

Achieving a comprehensive survey of cell types in the cerebellum poses some unique challenges. First, a large majority of the neurons are GCs, making it difficult to satisfactorily sample the rarer types. Second, for many of the morphologically and physiologically defined cell types—especially the interneuron populations—existing molecular characterization is extremely limited. Recent advances in single-cell RNA sequencing (scRNAseq) technology^2–4^ have increased the throughput of profiling to now enable the systematic identification of cell types and states throughout the central nervous system^5–8^. Several recent studies have harnessed such techniques to examine cell populations in the mouse cerebellum across developmental stages^9–11^, but none has yet surveyed cell types in the adult.

## Transcriptional definition of cerebellar cell types

We developed a pipeline for high-throughput single-nucleus RNA-seq (snRNA-seq) with exceptional transcript capture efficiency and nuclei yield, as well as consistent performance across regions of the adult mouse brain^12^ (Methods). To comprehensively sample cell types in the mouse cerebellum, we dissected and isolated nuclei from 16 different lobules, across both female and male replicates (Fig. 1a, Methods, Extended Data Fig. 1a). We recovered 780,553 nuclei profiles with a median transcript capture of 2862 unique molecular identifiers (UMIs) per profile (Extended Data Fig 1b, c), including 530,063 profiles from male donors, and 250,490 profiles from female donors, with minimal inter-individual batch effects (Extended Data Fig. 1d,e).

**Figure 1:**
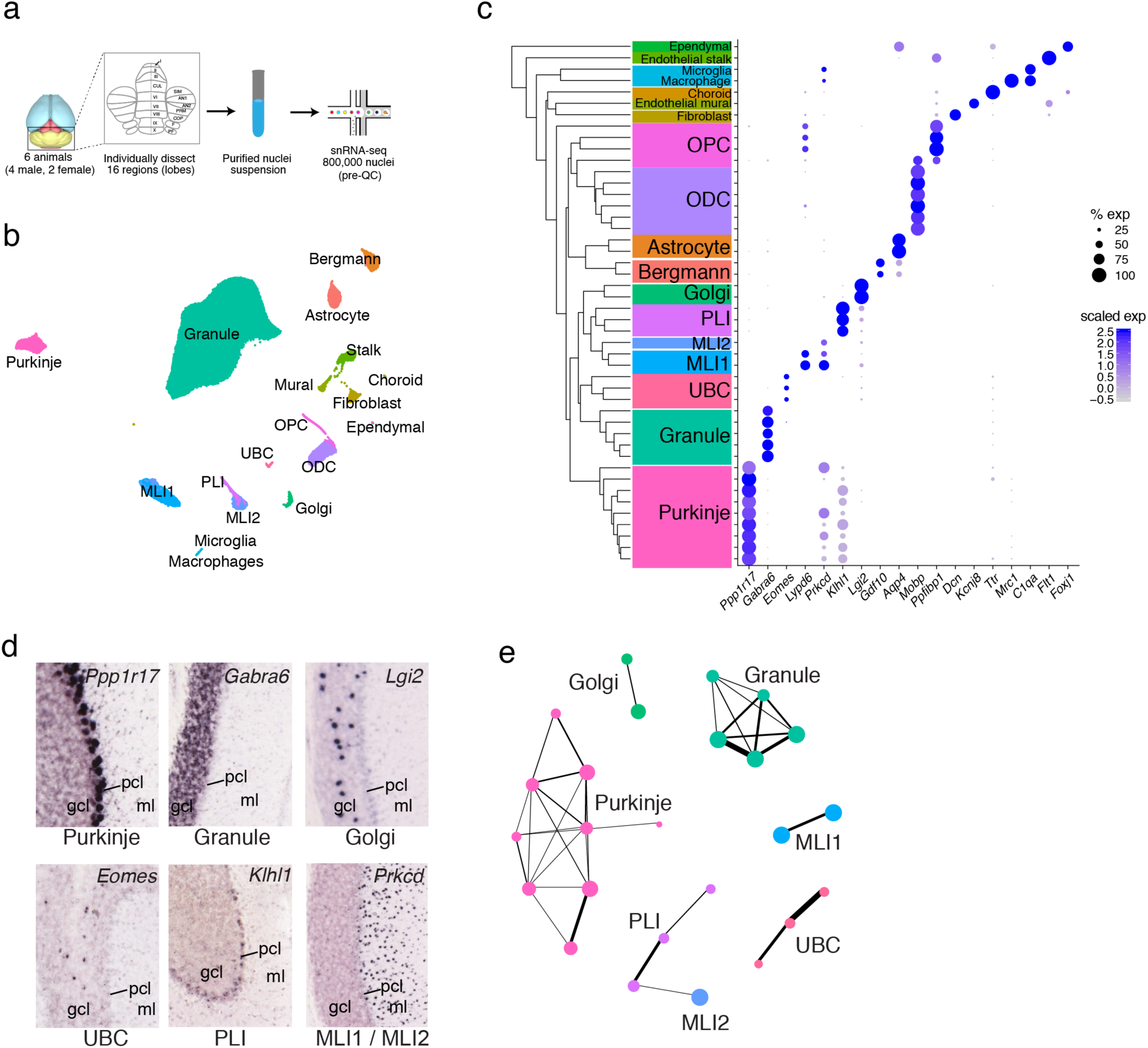
Comprehensive transcriptional profiling of cell types across the mouse cerebellum. **a**, Experimental design with lobe-based sampling and profiling. **b**, UMAP visualization of 611,034 nuclei (after profile QC, see Methods), colored by cell type identity. **c**, Dendrogram indicating hierarchical relationships between cell subtypes (left) with a paired dot plot (right) of scaled expression of selected marker genes for cell type identity. Background text colors correspond to cell types as in (**b**). **d**, ABA expression staining for selected gene markers of canonical interneuron populations, indicating cerebellar layer localization. **e**, Neighbor-based connectivity (see Methods) visualized for all neuronal clusters, in force-directed layout. Thickness of connecting lines represents magnitude of connectivity. OPC, oligodendrocyte precursor cell; ODC, oligodendrocyte; PLI, purkinje layer interneuron; MLI, molecular layer interneuron; UBC, unipolar brush cell; *gcl*, granule cell layer; *pcl*, purkinje cell layer; *ml*, molecular layer.

To discover cell types, we used a clustering strategy we developed previously^13^ (Methods) to partition 611,034 high-quality profiles into 46 clusters. We estimate that with this number of profiles, we can expect to reasonably sample even extremely rare cell types (prevalence of 0.15%) with a probability of greater than 90%, suggesting we captured the great majority of transcriptional variation within the cerebellum (Extended Data Fig. 1f).

We assigned each cluster to one of 18 known cell type identities based upon expression of specific molecular markers known to correlate with defining morphological, histological, and/or functional features (Fig. 1b-c, Extended Data Table 1). These annotations were also corroborated by the expected layer-specific localizations of marker genes in the Allen Brain Atlas (ABA) (Fig. 1d). Several types contained multiple clusters, suggesting additional heterogeneity within those populations (Extended Data Fig. 2a-n, Supplementary Table 1). To compare the discreteness of these subsets within different types, we developed a graph-based metric (Methods) of connectivity within cluster pairs. Applying this metric revealed higher connectivity amongst individual clusters of certain neuron cell types (Fig. 1e), suggesting that some cluster distinctions vary more subtly and continuously than others.

## Characterization of spatial variation and patterning in neuronal and glial cell types

To quantify regional specialization of cell types, we examined how our clusters distributed proportionally across each lobule. We found that eight of our nine PC clusters, as well as several GC clusters and one Bergmann glial cluster, showed the most significantly divergent lobule compositions (Pearson’s chi squared, FDR < 0.001, see Methods) and exhibited greater than two-fold enrichment in at least one lobule (Fig. 2a). There was high concordance in regional composition of each of these types across replicates, suggesting consistent spatial enrichment patterns (Extended Data Fig. 3a).

**Figure 2:**
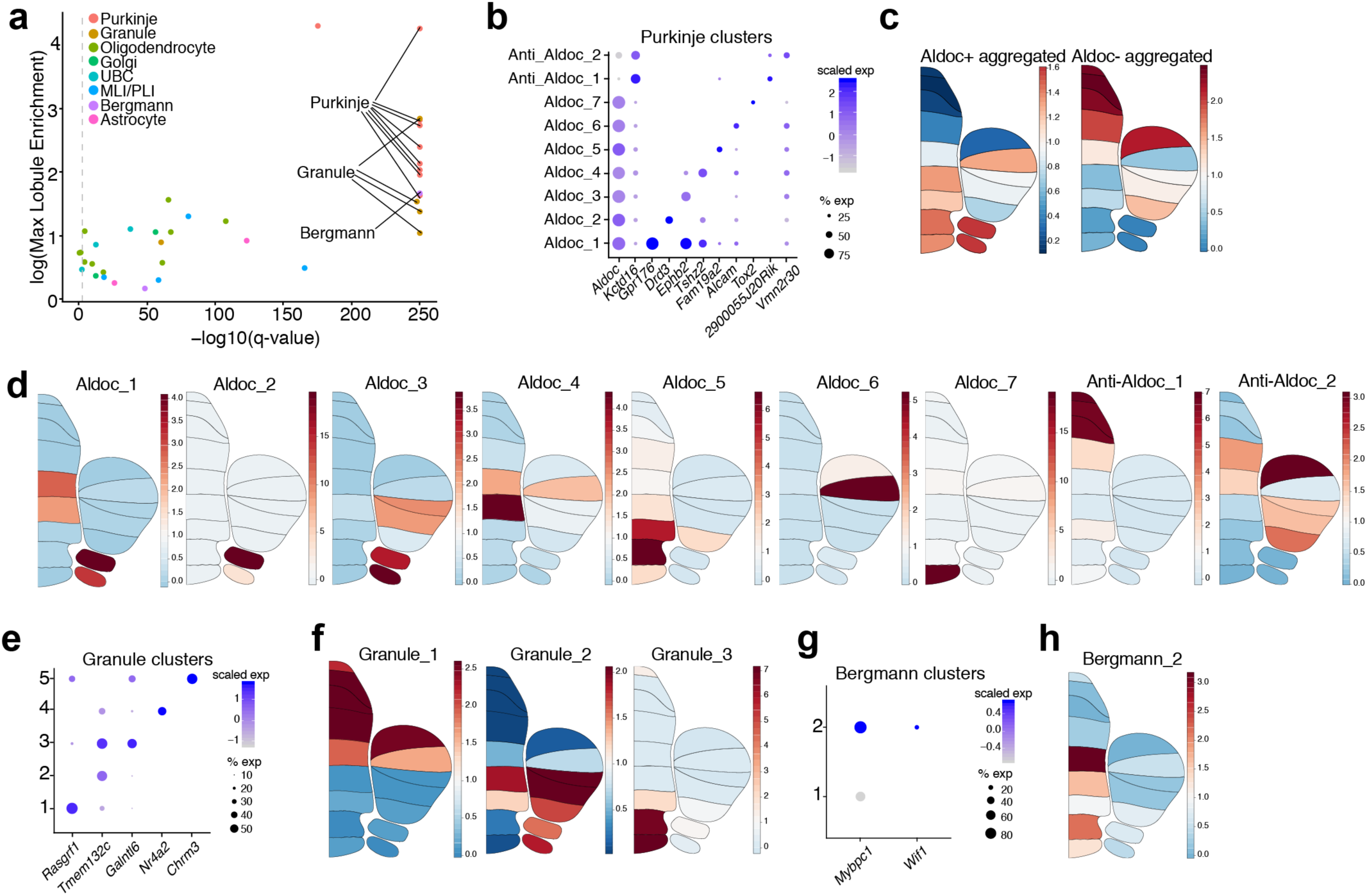
Characterization of spatial variation and patterning in neuronal and glial cell types. **a**, Scatter plot indicating neuronal and glial clusters which have lobule enrichment (LE) patterns significantly different from the cell type population as a whole (Pearson’s chi-squared, FDR < 0.001 indicated by dashed line). x axis shows -log10 transformed q values; y axis shows log2 transformed maximum lobule enrichment across all 16 lobules (see Methods). Genes with high correlation (Pearson correlation coefficient > 0.85) in LE values between replicate sets and max LE > 2 are labeled (see Methods, Extended Data Fig. 3a). **b**, Dot plot of scaled expression of selected gene markers for PC clusters. **c**, Regional enrichment plots indicating average lobule enrichment for aggregated *Aldoc*-positive PC subtypes (left) and aggregated *Aldoc*-negative subtypes (right). **d**, Regional enrichment plots indicating lobule enrichment for PC clusters. **e**, Dot plot of scaled expression of selected gene markers for GC clusters. **f**, Regional enrichment plots indicating lobule enrichment for three spatially significant GC clusters. **g**, Dot plot of scaled expression of selected gene markers for Bergman glial clusters. **h**, Regional enrichment plot indicating lobule enrichment for the *Mybpc1/Wif1+* Bergmann glial cluster.

The 9 PC clusters could be divided into two main groups, based upon their expression of *Aldoc*, which defines parasagittal striping of Purkinje neurons across the cerebellum^1^. Seven of the nine PC clusters were *Aldoc*+, indicating greater specialization in this population compared with the *Aldoc*-PCs. Combinatorial expression of *Aldoc* and at least one subtype-specific marker fully identified the Purkinje clusters (Fig. 2b). These *Aldoc*+ and *Aldoc-* groups showed a regional enrichment pattern consistent with the known paths of parasagittal stripes across individual lobules (Fig. 2c). In characterizing the spatial variation of the PC subtypes, we found some with spatial patterns we recently identified using Slide-seq technology (Aldoc_5 and Aldoc_7, marked by *Tox2* and *Gpr176* respectively)^14^, as well as several undescribed subtypes and patterns (Fig. 2b,d; Extended Data Fig. 3b). The majority of this Purkinje diversity was concentrated in the posterior cerebellum, particularly the uvula and nodulus, consistent with these regions showing greater diversity in function and connectivity^15,16^.

We also observed regional specialization in excitatory interneurons and Bergmann glia. Among the 5 GC subtypes (Fig. 2e), three had significant and cohesive spatial enrichment patterns (subtypes 1, 2, and 3, Fig. 2f, Extended Data Fig. 3c). In addition, consistent with prior work^17^, the UBCs were highly enriched in the posterior lobules (Extended Data Fig. 3d). Finally, we identified a Bergmann glial subtype expressing the marker genes *Mybpc1*^*14*^ and *Wif1* (Fig. 2g), with high enrichment in lobe VI and the nodulus (Fig. 2h, Extended Data Fig. 3e). The regional specialization of interneuron and glial populations stands in contrast to the cerebral cortex, where molecular heterogeneity across regions is largely limited to projection neurons^5,7^.

## Computational analysis of continuous and discrete subtype variation in unipolar brush cells and other neuronal cell types

Molecularly defined cell populations can be highly discrete—such as, in cerebral cortex, the distinctions between chandelier and basket interneuron types^6^—or they can vary more continuously, such as the cross-regional differences amongst principal cells of striatum^7,18^ and cortex^5,7^. The cerebellum is known to contain several canonical cell types that exist as morphological and functional continua, such as the basket and stellate interneurons of the molecular layer^19^. To examine continuous features of molecular variation in greater detail within interneuron types, we created a metric to quantify the continuity of gene expression. Briefly, we fit a logistic curve for expression of each gene along the dominant expression trajectory^20^, extracting the curve’s maximum slope (*m*) (Methods, Fig. 3a). Intuitively, we expect *m* values to be smaller for genes representative of a more continuous gradient of molecular expression (Fig. 3a).

**Figure 3:**
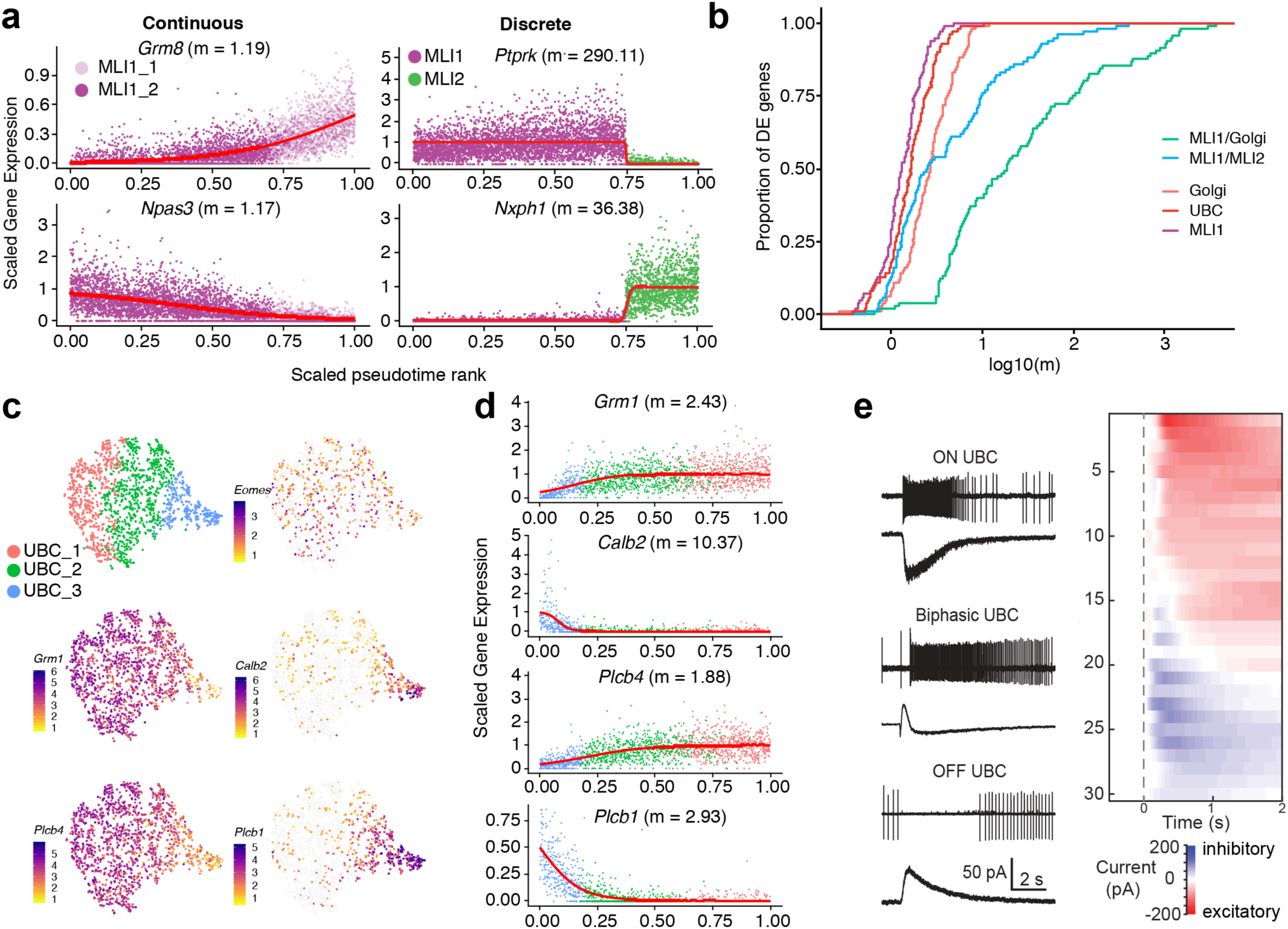
Cross-cluster continuity among select neuronal populations, including unipolar brush cells. **a**, Pseudotime ordered gene expression (Methods) for two of the top differentially expressed genes between two clusters within the MLI1 type (left), and between two cell types (right, MLI1 and MLI2 cell types). Curves indicate logistic fits estimated via nonlinear least squares; maximum slope values (*m*) indicated. Differences in magnitude of *m* values correspond well with visually distinctive molecular continuity versus discreteness. **b**, Empirical cumulative distributions of *m* values (curves fit as in (**a**)) for top differentially expressed genes among aggregated combinations of MLI, GoC and UBC clusters. **c**, t-SNE visualizations of UBCs (n = 1613), colored by cluster assignment (top left), and log-normalized expression of canonical gene markers for all UBCs (*Eomes*), ON UBCs (*Grm1, Plcb4*), and OFF UBCs (*Calb2, Plcb1*). **d**, Rank ordered gene expression for canonical markers associated with ON UBCs (*Grm1, Plcb4*) and OFF UBCs (*Calb2, Plcb1*). Curves indicate logistic fits estimated as in (**a**); maximum slope values (*m*) indicated. **e**, Cell-attached recordings of spiking responses and whole cell recordings of currents evoked by brief glutamate puffs in ON (top), biphasic (middle) and OFF (bottom) UBCs. Heatmap of the currents recorded in all UBCs, sorted by the magnitude and time course of charge transfer (left).

Comparing *m* values across 100 highly variable genes within the UBC, GoC, and MLI populations suggested that in UBCs, many genes showed continuous variation (Fig. 3b), consistent with the high degree of connectivity we observed across the three UBC clusters (Fig. 1e). Previously, UBCs were classified into discrete ON and OFF types, based upon their differential electrophysiological responses to mossy fiber input^21,22^. By contrast, our cluster analysis suggested that these ON and OFF populations defined extreme ends of a more continuous axis of variation. Indeed, within the intermediate cluster flanking the more discretely defined ON and OFF clusters, we observed joint expression of markers associated with OFF (*Calb2, Plcb1*) and ON (*Grm1, Plcb4*) UBCs (Fig. 3c,d).

We performed electrophysiological studies to determine whether the functional properties of UBCs reflect the diversity of molecular properties. We pressure-applied glutamate and measured the spiking responses of UBCs with on-cell recordings, and then broke into the cell to measure glutamate-evoked currents (Methods). In some cells, glutamate rapidly and transiently increased spiking and evoked a long-lasting inward current (Fig. 3e, top left). For other cells, glutamate transiently suppressed spontaneous firing and evoked an outward current (Fig. 3e, bottom left). Many UBCs, however, had more complex, mixed responses to glutamate; we refer to these as “biphasic” cells. In one cell, for example, glutamate evoked a delayed increase in firing, caused by an initial outward current followed by a longer lasting inward current (Fig. 3e, middle left). A summary of the glutamate evoked currents (Fig. 3e, right) suggests that the graded nature of the molecular properties of UBCs may lead to graded electrical response properties.

## Molecular layer interneurons are composed of two molecularly and functionally discrete types

MLIs are spontaneously active interneurons that inhibit PCs as well as other MLIs. MLIs are canonically subdivided into stellate cells located in the outer third of the ML, and basket cells located in the inner third of the molecular layer that synapse onto PC somata and form specialized contacts known as pinceaus, which ephaptically inhibit PCs. Many MLIs, particularly those in the middle of the ML, share morphological features with both basket and stellate cells^19^. Thus, MLIs are thought to represent a single functional and morphological continuum.

Our clustering analysis of MLIs and PLIs, by contrast, identified two discrete populations of MLIs. The first population, “MLI1”, uniformly expressed *Lypd6, Sorcs3*, and *Ptprk* (Fig. 1b,4a). The second population, “MLI2,” was highly molecularly distinct from MLI1, expressing numerous markers also found in PLIs, such as *Nxph1* and *Cdh22* (Fig. 4a). A cross-species analysis^13^ with profiles obtained from postmortem human cerebellum demonstrated that the MLI1 and MLI2 populations are both evolutionarily conserved (Fig. 4b). Single molecule FISH (smFISH) experiments with *Sorcs3* and *Nxph1* revealed the markers to be entirely mutually exclusive (Fig. 4c,d). Furthermore, both MLI1s and MLI2s were present throughout the entire molecular layer, indicating that the MLI1/MLI2 distinction does not correspond to the canonical basket/stellate distinction (Fig. 4c,d).

**Figure 4:**
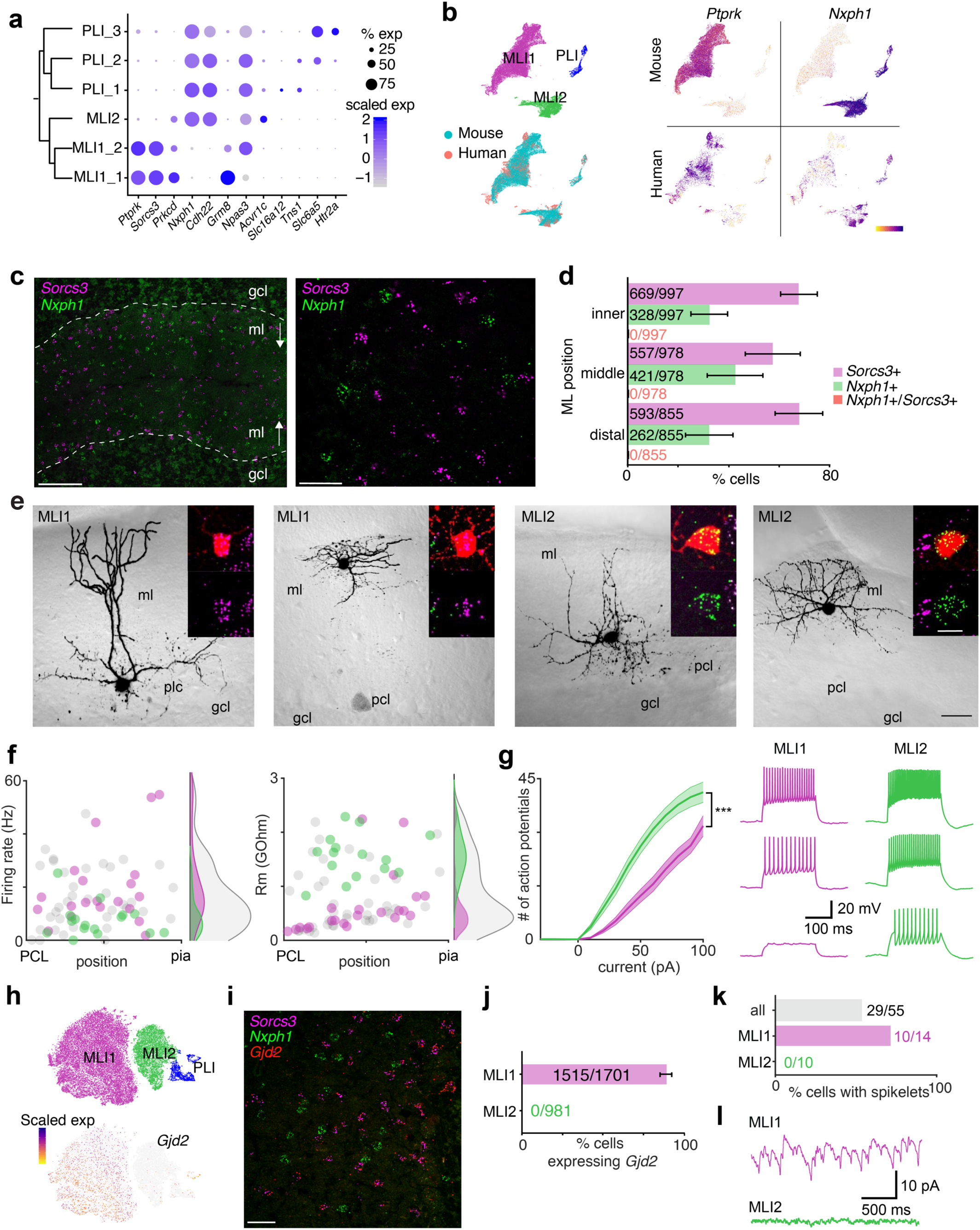
Two molecularly and electrophysiologically distinct populations of MLIs. **a**, Dendrogram of gene expression relationships amongst MLI and PLI populations (left), paired with dot plot (right) of selected gene markers. Dendrogram was computed as in Fig. 1b. **b**, t-SNE visualizations of cell types jointly identified by LIGER ^13^ across mouse and postmortem human nuclei, colored by cell type (top left) and species (bottom left). Middle and right, expression of MLI1 and MLI2 markers *Ptprk* and *Nxph1* respectively, subsetted by species (mouse top row, human bottom row). **c**, smFISH expression of *Sorcs3* (purple) and *Nxph1* (green) at low magnification (left, scale bar 100 μm) and higher magnification (right, scale bar = 20 μm). Dotted line indicates the location of the purkinje layer. **d**, Bar graph quantifying the percentages of *Sorcs3*+, *Nxph1*+ and *Sorcs3+*/*Nxph1*+ cells in the inner third, middle third, and distal third of the molecular layer, across 16 slides (total cells counted indicated). Error bars represent standard deviations. **e**, Four 2-photon images of representative basket and stellate-like MLI1 and MLI2 neurons in cerebellar slice. Insets denote confocal images of the fluorescent fill of the cell body (red) with the smFISH signals superimposed (top), and the smFISH signal only (bottom; *Sorcs3*, purple; *Nxph1*, green). Scale bars represent 30 mm (black) and 10 mm (white). **f**, Scatter plot of firing rate (left) and membrane resistance Rm (right) of molecularly identified MLI1 (purple circles) and MLI2 neurons (green circles), as well as MLIs whose molecular identity was not ascertained (gray circles). The corresponding distributions are shown to the right of the scatter plots. **g**, Left, Mean input-output curves of MLI1 (purple line) and MLI2 neurons (green line). Shaded area denotes standard error of the mean (SEM) values. Right, representative traces of MLI1 (purple) and MLI2 (green) for 20, 40 and 60 pA current injections. **h**, t-SNE visualization of expression of *Gjd2* across MLI and PLI cell types. **i**, Representative image of smFISH expression of *Sorcs3* (purple), *Nxph1* (green), and *Gjd2* (red), in the molecular layer (scale bar = 20 μm). **j**, Bar graph showing percentages of MLI1s and MLI2s expressing *Gjd2* in the molecular layer, across 16 slides (total cells counted indicated). Error bars represent standard deviations. **k**, Bar graph showing the percentages of all cells (grey bar), MLI1s (purple bar) and ML2s (green bar) in which spikelets were observed. **l**, Example voltage clamp recordings in the presence of synaptic blockers show spikelets in an MLI1 (top, purple trace), while MLI2 is devoid of spikelets (bottom, green trace). ml, molecular layer; pcl, purkinje cell layer; gcl, granule cell layer. Firing rates, Rm, input/output curves and numbers of cells with spikelets were all significantly different for MLI1 and MLI2 (Extended Data Table 2).

To better understand the morphological, physiological, and molecular characteristics of the MLI populations, we developed a pipeline to record from individual MLIs in brain slices, image their morphologies, and then ascertain their molecular MLI1/MLI2 identities by smFISH (Methods; Fig. 4e). Consistent with the marker analysis (Fig. 4a), MLI1s had a stellate morphology in the distal third of the ML, whereas MLI1s located near the PC layer had a basket morphology, with contacts near PC initial segments (Fig. 4e; Extended Data Fig. 4). We next examined whether MLI2s, which did not show any obvious systematic molecular heterogeneity, had graded morphological properties. MLI2s in the distal third of the ML also had stellate cell morphology, whereas MLI2s near the PC layer had a distinct morphology and appeared to form synapses preferentially near the PC layer (Extended Data Fig. 4). Although further studies are needed to determine if MLI2s form pinceaus, it is clear that both MLI1 and MLI2 showed a similar gradient in their morphological properties.

The electrical characteristics of MLI1s and MLI2s showed numerous distinctions. The average spontaneous firing rate was significantly higher for MLI1s than MLI2s (Mann-Whitney test, p=0.0015) (Fig. 4f). The membrane resistance (Rm) of MLI1s was lower than that of MLI2s (Fig. 4f). We also found that MLI2s were more excitable than MLI1s (Fig. 4g), and displayed a stronger hyperpolarization-activated current (Extended Data Fig. 5).

MLIs are known to be electrically coupled via gap junctions^23^, but it is not clear if this is true for both MLI1s and MLI2s. In the cerebral cortex and some other brain regions, interneurons often electrically couple selectively to neurons of the same type, but not other types^24,25^. We therefore examined whether this is also true for MLI1s and MLI2s. Expression of *Gjd2*, the gene encoding the dominant gap junction protein in MLIs^26^, was found in MLI1s but not MLI2s, both in our single-nucleus data (Fig. 4h), and by smFISH (Fig. 4i,j), suggesting potential differences in electrical coupling. Action potentials in coupled MLIs produce small depolarizations known as spikelets that are thought to promote synchronous activity between MLIs^23^. We therefore investigated whether spikelets are present in MLI1s and absent in MLI2s. Consistent with the gene expression profile, we observed spikelets in 71% of MLI1s and never in MLI2s (Fig. 4k,l; p < 10^−3^, Fisher’s exact test). These findings suggest that most MLI1s are gap junction coupled to other MLI1s, while MLI2s show no electrical coupling to other MLIs.

## Conclusions

Here, we used high-throughput, region-specific transcriptome sampling to build a comprehensive taxonomy of cell types in the mouse cerebellum, and quantify spatial variation across individual regions. Our dataset is freely available to the neuroscience community^27,28^, facilitating functional characterization of these populations, many of which are entirely novel. One of the biggest challenges facing the comprehensive cell typing of the brain is the correspondence problem^29^: how to integrate definitions of cell types based on many modalities of measurement used to characterize brain cells. We were surprised to find that the cerebellar MLIs—one of the first sets of neurons to be characterized over 130 years ago^30^—are in fact composed of two molecularly and physiologically discrete populations, that each themselves show a similar morphological continuum along the depth axis of the ML. Only through joint characterization of gene expression, morphology, and physiology in the same cell were we able to clarify MLI heterogeneity, underscoring the importance of multi-modal studies for brain cell typing. As comprehensive cell typing proceeds across other brain regions, we expect the emergence of similar basic discoveries that challenge and extend our understanding of cellular specialization in the nervous system.

## Acknowledgments

We would like to thank Aleksandrina Goeva for helpful discussion. This work was supported by the Stanley Center for Psychiatric Research, and NIH/NIMH Brain Grant 1U19MH114821 to E.Z.M.

## Author Contributions

V.K. performed analyses, with help from C.M. C.M. generated the single-nucleus profiles and performed the clustering analysis. C.G. and W.R. conceived UBC physiology experiments, which were performed by C.G. W.R., T.O. and S.R. conceived MLI electrophysiology/imaging experiments, which were performed and analyzed by T.O. and S.R.. C.V. performed dissections. N.N., C.V., and C.M. performed the smFISH experiments. E.Z.M. conceived the molecular study, with help from A.R. E.Z.M. and V.K. wrote the paper with contributions from all authors.

## Methods

### Animals

Nuclei suspensions for mouse cerebellum profiles were generated from a total of 2 adult female and 4 adult male mice (60 days old; C57BL/6J, Jackson Labs). Animals were group-housed with a 12-hour light-dark schedule and allowed to acclimate to their housing environment for two weeks post-arrival. All experiments were approved by and in accordance with Broad IACUC protocol number 012-09-16.

### Brain preparation

At 60 days of age, C57BL/6J mice were anesthetized by administration of isoflurane in a gas chamber flowing 3% isoflurane for 1 minute. Anesthesia was confirmed by checking for a negative tail pinch response. Animals were moved to a dissection tray and anesthesia was prolonged via a nose cone flowing 3% isoflurane for the duration of the procedure. Transcardial perfusions were performed with ice cold pH 7.4 HEPES buffer containing 110 mM NaCl, 10 mM HEPES, 25 mM glucose, 75 mM sucrose, 7.5 mM MgCl2, and 2.5 mM KCl to remove blood from the brain and other organs sampled. The brain was removed and frozen for 3 minutes in liquid nitrogen vapor and moved to -80C for long term storage. A detailed protocol is available at protocols.io (dx.doi.org/10.17504/protocols.io.bcbrism6).

### Generation of cerebellar nuclei profiles

Frozen mouse brains were securely mounted by the frontal cortex onto cryostat chucks with OCT embedding compound such that the entire posterior half including the cerebellum and brainstem were left exposed and thermally unperturbed. Dissection of each cerebellar vermal and cortical lobe was performed by hand in the cryostat using an ophthalmic microscalpel (Feather safety Razor #P-715) precooled to -20°C and donning 4x surgical loupes. Each excised tissue dissectate was placed into a pre-cooled 0.25 ml PCR tube using pre-cooled forceps and stored at -80°C. Nuclei were extracted from this frozen tissue using gentle, detergent-based dissociation, according to a protocol available at protocols.io (dx.doi.org/10.17504/protocols.io.bck6iuze) adapted from one generously provided by the McCarroll lab (Harvard Medical School), and loaded into the 10x Chromium V3 system. Reverse transcription and library generation were performed according to the manufacturer’s protocol.

### Floating Slice Hybridization Chain reaction (HCR) on acute slices

Acute cerebellar slices containing Alexa 594-filled patched cells were fixed as described and stored in 70% ethanol at 4°C until HCR. They were then subjected to a “floating slice HCR’’ protocol whereby the recorded cells could be simultaneously re-imaged in conjunction with HCR expression analysis *in situ* and cataloged as to their positions in the cerebellum. A detailed protocol (dx.doi.org/10.17504/protocols.io.bck7iuzn) was performed using the following HCR probes and matching hairpins purchased from Molecular Instruments, Inc. (Los Angeles, CA): Glutamate metabotropic receptor 8 (*Grm8*) lot number PRC005, Connexin 36 (*Gjd2*) lot number PRD854, Cadherin22 (*Cdh22*) lot number PRC011, Neurexophilin 1 (*Nxph1*) lot number PRC675 and PRC466 and Sortilin related VPS10 domain containing receptor 3 (*Sorcs3*) lot number PRC004. Amplification hairpins used were type B1, B2 and B3 in 488nm, 647nm and 546nm respectively.

### Patch fill and HCR co-imaging

Following floating slice HCR, slices were mounted between no.1 coverslips with antifade compound (ProLong Glass, invitrogen) and images were collected on an Andor CSU-X spinning disk confocal system coupled to a Nikon Eclipse Ti microscope equipped with an Andor iKon-M camera. The images were acquired with an oil immersion objective at 60x. The Alexa 594 patched cell backfill channel (561nm) plus associated HCR probe/hairpin channels (488nm and 647nm) were projected through a 10-20 micron thick z-series so that an unambiguous determination of the association between the patch-filled cell and its HCR gene expression could be made. Images were processed using Nikon NIS Elements 4.4 and ImageJ.

### Human brain and nuclei processing

Human donor tissue was supplied by the Human Brain and Spinal Fluid Resource Center at UCLA, through the NIH NeuroBioBank. This work was determined by the Office of Research Subjects Protection at the Broad Institute not to meet the definition of human subjects research (project ID NHSR-4235).

Nuclei suspensions from human cerebellum were generated from two neuropathologically normal control cases, one female tissue donor, aged 35 and one male tissue donor, aged 36. These fresh frozen tissues had postmortem intervals of 12 and 13.5 hours respectively, and were provided as whole cerebella cut into 4 coronal slabs. A sub-dissection of frozen cerebellar lobules was performed on dry ice just prior to 10x processing and nuclei were extracted from this frozen tissue using gentle, detergent-based dissociation, according to a protocol available at protocols.io (dx.doi.org/10.17504/protocols.io.bck6iuze).

### Electrophysiology experiments

Acute parasagittal slices were prepared at 240 μm thickness from wild type mice aged P30–50. Mice were anesthetized with an IP injection of ketamine (10 mg/kg), perfused transcardially with an ice-cold solution containing (in mM): 110 CholineCl, 7 MgCl_2_, 2.5 KCl, 1.25 NaH_2_PO_4_, 0.5 CaCl_2_, 25 Glucose, 11.5 Na-ascorbate, 3 Na-pyruvate, 25 NaHCO_3_, 0.003 (R)-CPP, equilibrated with 95% O_2_ and 5% CO_2_. Slices were cut in the same solution and were then transferred to artificial cerebrospinal fluid (ACSF) containing (in mM) 125 NaCl, 26 NaHCO_3_, 1.25 NaH_2_PO_4_, 2.5 KCl, 1 MgCl_2_, 1.5 CaCl_2_, and 25 glucose equilibrated with 95% O_2_ and 5% CO_2_ at ∼34 °C for 30 min. Slices were then kept at room temperature until recording.

All UBC recordings were done at 34 to 36 °C with (in µM) 2 R-CPP, 5 NBQX, 1 strychnine, 10 SR95531 (gabazine), 1.5 CGP in the bath to isolate metabotropic currents. Loose cell-attached recordings were made with ACSF-filled patch pipettes of 3-5 MΩ resistance. Whole-cell voltage-clamp recordings were performed while holding the cell at -70 mV with an internal containing (in mM): 140 KCl, 4 NaCl, 0.5 CaCl_2_, 10 HEPES, 4 MgATP, 0.3 NaGTP, 5 EGTA 5, and 2 QX-314, pH adjusted to 7.2 with KOH. Brief puffs of glutamate (1 mM conc. for 50 ms at 5 psi) were delivered using a Picospritzer™ II (General Valve Corp., Fairfield, NJ, USA) in both cell-attached and whole-cell configuration to assure consistent responses. The heatmap of current traces from all cells are sorted by the score over the first principal axis after singular value decomposition (SVD) of recordings over all cells.

MLI recordings were performed at ∼32 °C with an internal solution containing (in mM) 150 K-gluconate, 3 KCl, 10 HEPES, 3 MgATP, 0.5 GTP, 5 phosphocreatine-tris_2_, and 5 phosphocreatine-Na_2_, 2 mg/ml biocytin and 0.1 Alexa 594 (pH adjusted to 7.2 with KOH, osmolality adjusted to 310 mOsm/kg). Visually guided whole-cell recordings were obtained with patch pipettes of ∼4 MΩ resistance pulled from borosilicate capillary glass (BF150-86-10, Sutter Instrument, Novato, CA). Electrophysiology data was acquired using a Multiclamp 700B amplifier (Axon Instruments), digitized at 20 kHz and filtered at 4 kHz. For isolating spikelets in MLI recordings, cells were held at -65 mV in voltage clamp and the following receptor antagonists were added to the solution (in µM) to block synaptic currents: 2 R-CPP, 5 NBQX, 1 strychnine, 10 SR95531 (gabazine), 1.5 CGP. All drugs were purchased from Abcam (Cambridge, MA) and Tocris (Bristol, UK). To obtain an input-output curve, MLIs were injected with a constant hyperpolarizing current to hold them at ∼60-65 mV, and 250 ms long current steps ranging from -30 pA to +100 pA were injected in 10 pA increments. To activate the hyperpolarization-evoked current (I_h_), MLIs were held at -65 mV and a 30 pA hyperpolarizing current step of 500 ms duration was injected. The amplitude of I_h_ was calculated as the difference between the maximal current evoked by the hyperpolarizing current step and the average steady-state current at the end (480-500ms) of the current step. Capacitance and input resistance (Ri) were determined using a 10 pA, 50 ms hyperpolarizing current step. To prevent excessive dialysis and to ensure successful detection of mRNAs in the recorded cells, the total duration of recordings did not exceed 10 min. Acquisition and analysis of electrophysiological data were performed using custom routines written in MATLAB (Mathworks, Natick, MA), IgorPro (Wavemetrics, Lake Oswego, OR), or AxoGraphX. Data are reported as median ± interquartile range, and statistical analysis was carried out using the Mann-Whitney or Fisher’s exact test, as indicated. Statistical significance was assumed at p < 0.05.

To determine the presence of spikelets, peak detection was used to generate event-triggered average waveforms with thresholds based on the mean absolute deviation (MAD) of the raw trace. Spikelet recordings were scored for the presence of spikelets blind to the molecular identity of the cells. The analysis was restricted to cells recorded in the presence of synaptic blockers.

### Imaging and analysis

MLIs were filled with 100 μM Alexa-594 via patch pipette to visualize their morphology using 2-photon imaging. After completion of the electrophysiological recordings the patch electrode was retracted slowly and the cell resealed. We used a custom-built 2-photon laser-scanning microscope with a 40x, 0.8 numerical aperture (NA) objective (Olympus Optical, Tokyo, Japan) and a pulsed 2-photon laser (Chameleon or MIRA 900, Coherent, Santa Clara, CA, 800 nm excitation). DIC images were acquired at the end of each experiment and locations of each cell within the slice were recorded. 2-photon images were further processed in ImageJ.

### Tissue fixation of acute slices

After recording and imaging, cerebellar slices were transferred to a well-plate and submerged in 2-4% PFA in PBS (pH=7.4) and incubated overnight at 4 °C. Slices were then washed in PBS (3×5min) and then kept in 70% Ethanol in RNAse-free water until HCR was performed.

### Preprocessing of sequencing reads

Sequencing reads from mouse cerebellum experiments were demultiplexed and aligned to a mouse (mm10) premrna reference using CellRanger v3.0.2 with default settings. Digital gene expression matrices were generated with the CellRanger count function. Sequencing reads from human cerebellum experiments were demultiplexed and aligned to a human (hg19) premrna reference using the Drop-seq alignment workflow^2^, which was also used to generate the downstream digital gene expression matrices.

### Estimation of adequate rare cell type detection

To estimate the probability of sufficiently sampling rare cell types in the cerebellum as a function of total number of nuclei sampled, we used the approach proposed by the the Satija lab (https://satijalab.org/howmanycells), with the assumption of 10 very rare cell types, each with a prevalence of 0.15%. We derived this minimum based on the observed prevalences of the two rarest cell types we identified (OPC_4, Purkinje_Aldoc_2). We set 70 cells as the threshold for sufficient sampling, and calculated the overall probability as a negative binomial density:

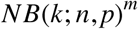

with k = 70, p = 0.0015, m = 10, and n representing the total number of cells sampled.

### Cell type clustering and annotation

After generation of digital gene expression matrices as described above, we filtered out nuclei with fewer than 500 UMIs. We then performed cell type annotation iteratively through a number of rounds of dimensionality reduction, clustering, and removal of putative doublets and cells with high mitochondrial expression. For the preliminary clustering step, we performed standard preprocessing (UMI normalization, highly variable gene selection, scaling) with Seurat v2.3.4 as described in Butler et al^31^. We used principal component analysis (PCA) with 30 components and Louvain community detection with resolution 0.1 to identify major clusters (resulting in 34 clusters). At this stage, we merged several clusters (primarily granule cell clusters) based on shared expression of canonical cell type markers, and removed one cluster whose top differentially expressed genes were mitochondrial (resulting in 10 clusters).

For subsequent rounds of cluster annotation within these major cell type clusters, we applied a variation of the LIGER workflow previously described^13^, using integrative non-negative matrix factorization (iNMF) to limit the effects of sample- and sex-specific gene expression. Briefly, we normalized each cell by the number of UMIs, selected highly variable genes^7^ and spatially variable genes (see next section), performed iNMF, and clustered using Louvain community detection (omitting the quantile alignment step). Clusters whose top differentially expressed genes indicated contamination from a different cell type or high expression of mitochondrial genes were removed during the annotation process, and not included in subsequent rounds of annotation. This iterative annotation process was repeated until no contaminating clusters were identified in a round of clustering. Differential expression analysis within rounds of annotation was performed with the Wilcoxon rank sum test using Seurat’s FindAllMarkers function. Final differential expression analysis across all 46 clusters was performed using the wilcoxauc function from the presto package^32^. A full set of parameters used in the LIGER annotation steps can be found in Extended Data Table 3.

For visualization as in Fig. 1B we merged all annotated high-profile nuclei and repeated preliminary preprocessing steps before performing UMAP on 25 principal components.

### Integrated analysis of mouse and human data

After generation of digital gene expression matrices for the human nuclei profiles, we filtered out nuclei with fewer than 500 UMIs. We then performed a preliminary round of cell type annotation using the standard LIGER workflow (integrating across batches) to identify the human molecular layer (ML) and Purkinje l ayer (PL) i nterneuron populations (based on the same markers as i n Extended Data Table 1). We repeated an iteration of the same workflow (with an additional quantileAlignSNF step) i n order to i dentify and remove putative doublet and contamination populations. Finally we performed an integrated LIGER analysis (integrating across species) using the remaining human ML/PL interneuron populations and the annotated MLI/PLI mouse populations.

### Spatially variable gene selection

To identify genes with high regional variance, we first computed the l og of the i ndex of dispersion (log variance-to-mean ratio, “logVMR”) for each gene, across each of the 16 lobular regions. Next, we simulated a Gaussian null distribution whose center was the logVMR mode, found by performing a kernel density estimation of the l ogVMRs (using the density function i n R, followed by the turnpoints function). The standard deviation of the Gaussian was computed by reflecting the values l ess than the mode across the center. Genes whose l ogVMRs were i n the upper tail with p < 0.01 (Benjamini-Hochberg adjusted) were ruled as spatially variable. For the GC and PC cluster analyses alpha thresholds were set to 0.001 and 0.002 respectively.

### Cluster regional composition test and lobule enrichment

To determine whether a cluster’s lobule composition differs significantly from the corresponding cell type lobule distribution, we used a multinomial test approximated by Pearson’s chi-squared test with k-1 degrees of freedom, where k was the total number of lobules sampled (16). The expected number of nuclei for a cluster *i* and lobule *j* was estimated as follows:

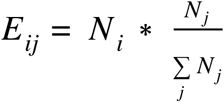

where *N*_*i*_ is the total number of nuclei in cluster *i* and *N*_*j*_ is the total number of nuclei in lobule *j* (across all clusters in the outer level cell type). The resulting p-values were FDR (Benjamini-Hochberg) adjusted using the p.adjust function in R.

Lobule enrichment scores for each cluster *i* and each lobule *j* were calculated according to:

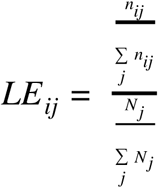

where *n*_*ij*_ is the observed number of nuclei in cluster *i* and lobule *j*, and *N*_*j*_ is the total number of nuclei in lobule *j* (across all clusters in the outer level cell type).

To determine consistency of lobule enrichment scores across replicates in each region, we designated two sets of replicates by assigning nuclei from the most represented replicate in each region and cluster analysis to “Replicate 1” and nuclei from the second most represented replicate in each region to “Replicate 2”. This assignment was done since not all regions had representation from all individuals profiled, and some had representation from only two individuals. We calculated lobule enrichment scores for each cluster using each of the replicate sets separately; we then calculated the Pearson correlation between the two sets of lobule enrichment scores for each cluster. Intuitively, we would expect correlation to be high for clusters when lobule enrichment is biologically consistent.

### Cluster connectivity analysis

To quantify relationships between all pairs of the 25 neuronal clusters (Fig. 1e), we quantified the proportion of shared nearest neighbor (SNN) graph connectivity found between the cells of one cluster versus each other cluster. Specifically, we first performed principal component analysis on scaled values for the 7718 genes that were called as highly variable in any of the neuron clustering analyses. Using the first 120 PCs, we constructed an SNN as previously described^33^, using the BuildSNN function in Seurat. For a given cluster pair X and Y, the connectivity metric was defined as the sum of X’s nonzero edge values with cells in cluster Y, divided by all nonzero edge values for the cells in cluster X. Note that this metric is asymmetric, and thus the connectivity from Y to X will be a different value than the connectivity to X to Y. The weighted edges in Fig. 1e represent the mean of the two connectivity values (*e.g.* X to Y and Y to X).

The force directed layout of neuronal subtypes in Fig. 1e was generated using Scanpy v1.4.4 (with sc.pl.paga), using PCs calculated in Seurat and connectivity values as described above.

### Continuity of gene expression

To characterize molecular variation across cell types, we attempted to quantify the continuity of scaled gene expression across a given cell type pair, ordered by pseudotime rank (calculated using Monocle2). For each gene, we then fit a logistic curve to the scaled gene expression values and calculated the maximum slope (*m*) of the resulting curve, after normalizing for both the number of cells and dynamic range of the logistic fit. To limit computational complexity, we downsampled cell type pairs to 5000 total nuclei.

We fit curves and calculated *m* values for the most significantly differentially expressed genes across 5 cell type pairs (Fig. 3b). Differentially expressed genes were determined using Seurat’s FindMarkers function. We then plotted the cumulative distribution of m values for the top 100 genes for each cell type pair; genes were selected after ordering by absolute Spearman correlation between scaled gene expression and pseudotime rank.

## Data Availability

All processed data and annotations have been made freely available for download and visualization through an interactive Single Cell Portal study (https://singlecell.broadinstitute.org/single_cell/study/SCP795/). Raw and processed data that support the findings of this study have been deposited in GEO under accession number XXX and in at the Neuroscience Multi-omics (NeMO) Archive (https://nemoarchive.org/)

---

**Extended Data Figure 1:**
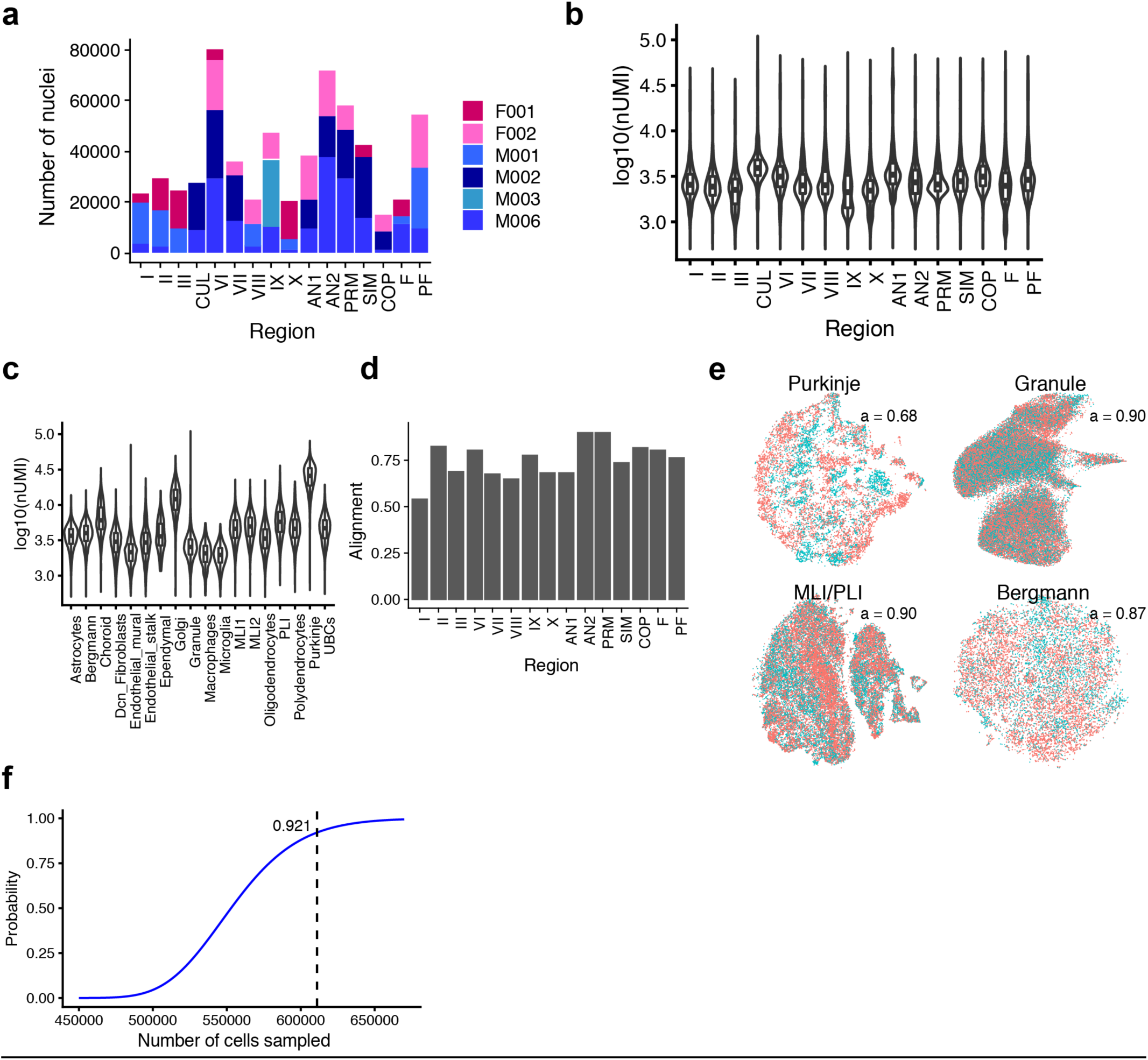
Summary and quality control analyses for nuclei sampling. **a**, Bar graph showing number of cells contributed by each individual per region across dataset of 611,034 nuclei (post-QC, 6 total individuals, 16 regions). **b**, Violin plot showing distribution of log10(nUMI) across the lobules profiled. **c**, Violin plot of log10(nUMI) per profile across the 18 cell types identified. The relative median values here are consistent with known differences in cell size; for example, Purkinje cells have the highest median number of UMIs. **d**, Bar graph of alignment scores (Methods) calculated across replicates for each lobule, after performing LIGER integration (across sex) (Methods) for each regional subset. Subsets sampled from the final set of 611,034 high-quality nuclei profiles. These analyses represent examples of expected replicate alignment when using the described pipeline. Note that lobule COP is excluded as it did not include representation from male and female replicates. **e**, Visualizations of representative cell type analyses, indicating high alignment across replicate sets (Granule is UMAP, all others t-SNE). These represent final analyses in the annotation process. Replicate sets were designated as in Methods (Cluster regional composition test and lobule enrichment). **f**, Plot indicating probability of sufficiently sampling 10 very rare populations (prevalence 0.15%) as a function of total number of cells profiled in experiment (probability estimated as in Methods). Number of high-quality nuclei profiled here (611,034) and corresponding probability are indicated.

**Extended Data Figure 2:**
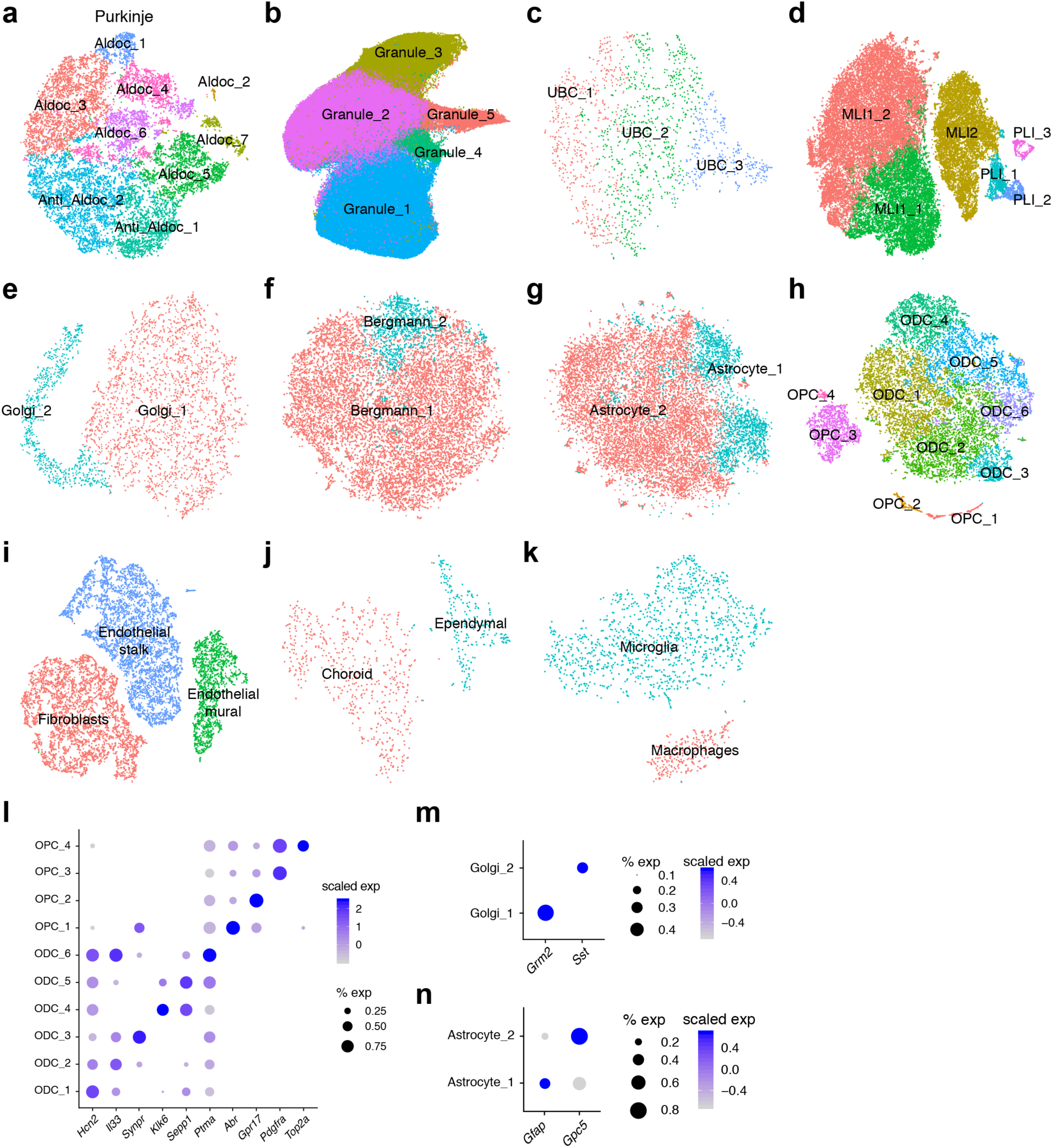
Characterization and annotation of cerebellar subtypes. **a** - **k**, Visualizations of all individual cell type analyses of purkinje (**a**), granule (**b**), UBC (**c**), MLI/PLI (**d**), and golgi (**e**) neurons, as well as bergmann (**f**), astrocyte (**g**), OPC/ODC (**h**), endothelial (**i**), choroid (**j**), and macrocytic (**k**) glial populations, labeled by cluster designations. Granule is UMAP, all others t-SNE. **l-n**, Dot plots of scaled expression of selected marker genes for three individual cell type analyses not displayed in the main figures: ODC/OPC (**l**), Golgi (**m**), and astrocyte (**n**).

**Extended Data Figure 3:**
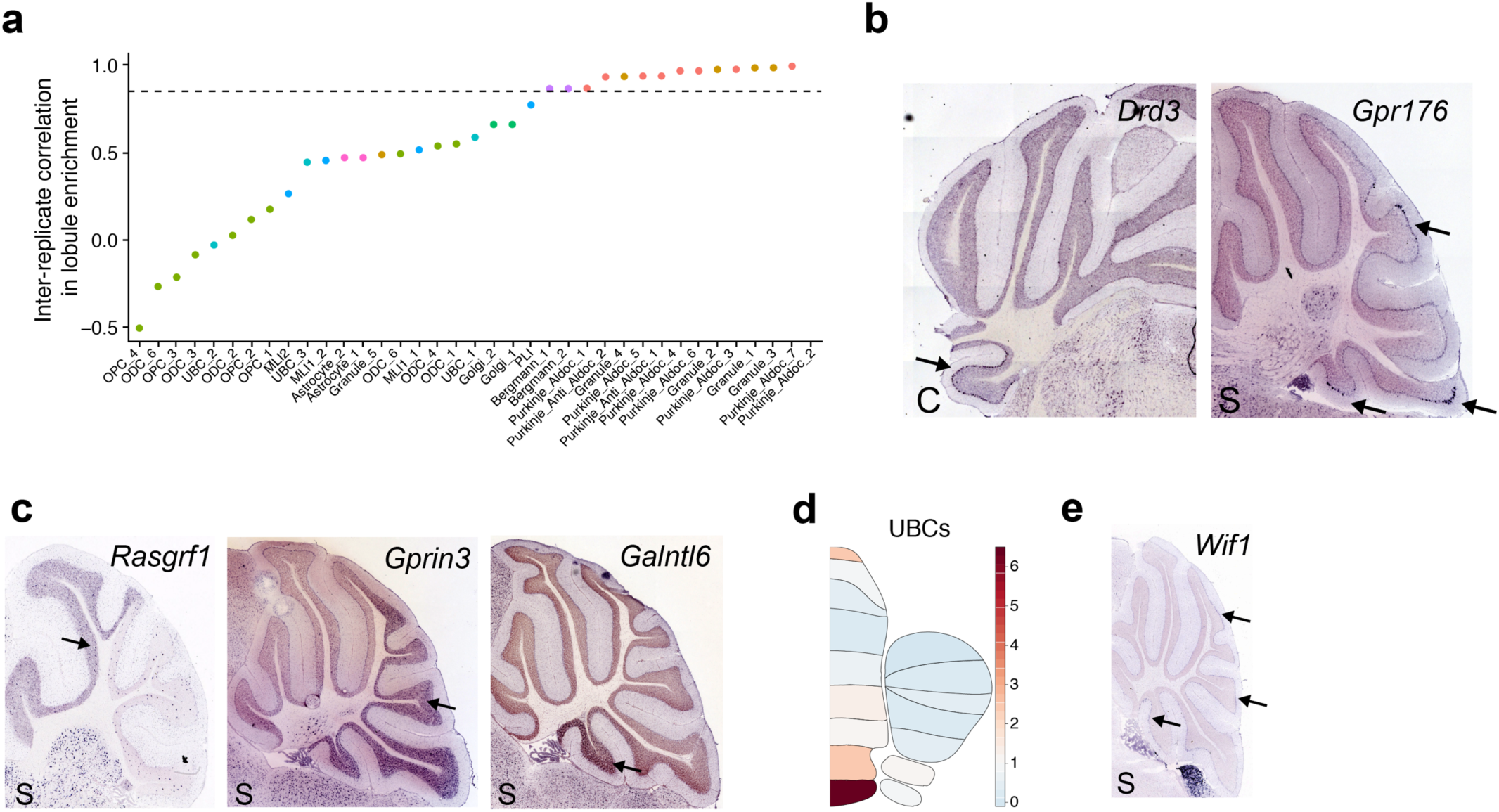
Additional analyses of spatial variation in neuronal and glial subtypes. **a**, Scatter plot of inter-replicate correlation (Pearson) for lobule enrichment scores calculated for replicate sets individually, across each cluster (clusters ordered by decreasing correlation). Two replicate sets were designated for each major cluster analysis by aggregating the individuals with the highest representation for each lobule into a single replicate (and similarly for the individuals with second highest representation). High inter-replicate correlation indicates consistent lobule enrichment for subtypes. Threshold of p = 0.85 is indicated. **b**, Allen Brain Atlas (AB) expression staining for two selected PC markers representing clusters with their respective lobule enrichments indicated; *Drd3* in the flocculus (Purkinje_Aldoc_2), and *Gpr176* in lobes VII, IX (uvula) and nodulus (X) (Purkinje_Aldoc_1). C indicates coronal section; S indicates sagittal section. **c**, ABA expression staining for three selected GC markers representing clusters with their respective lobule enrichments indicated; *Rasgrf1* in the anterior lobes (Granule_1), *Gprin3* in the posterior lobes (Granule_2), and *Galntl6* in the nodulus (Granule_3). **d**, Lobule enrichment plot indicating enrichment of UBCs in posterior lobules of the cerebellum, particularly lobes IX (uvula) and X (nodulus). Note that there is also slight enrichment in lobe VII, as suggested by previous studies. **e**, ABA expression staining for *Wif1* (a Bergmann_2 cluster marker), indicating expression enriched in lobe VI, lobe X (nodulus), and lobe VIII.

**Extended Data Figure 4:**
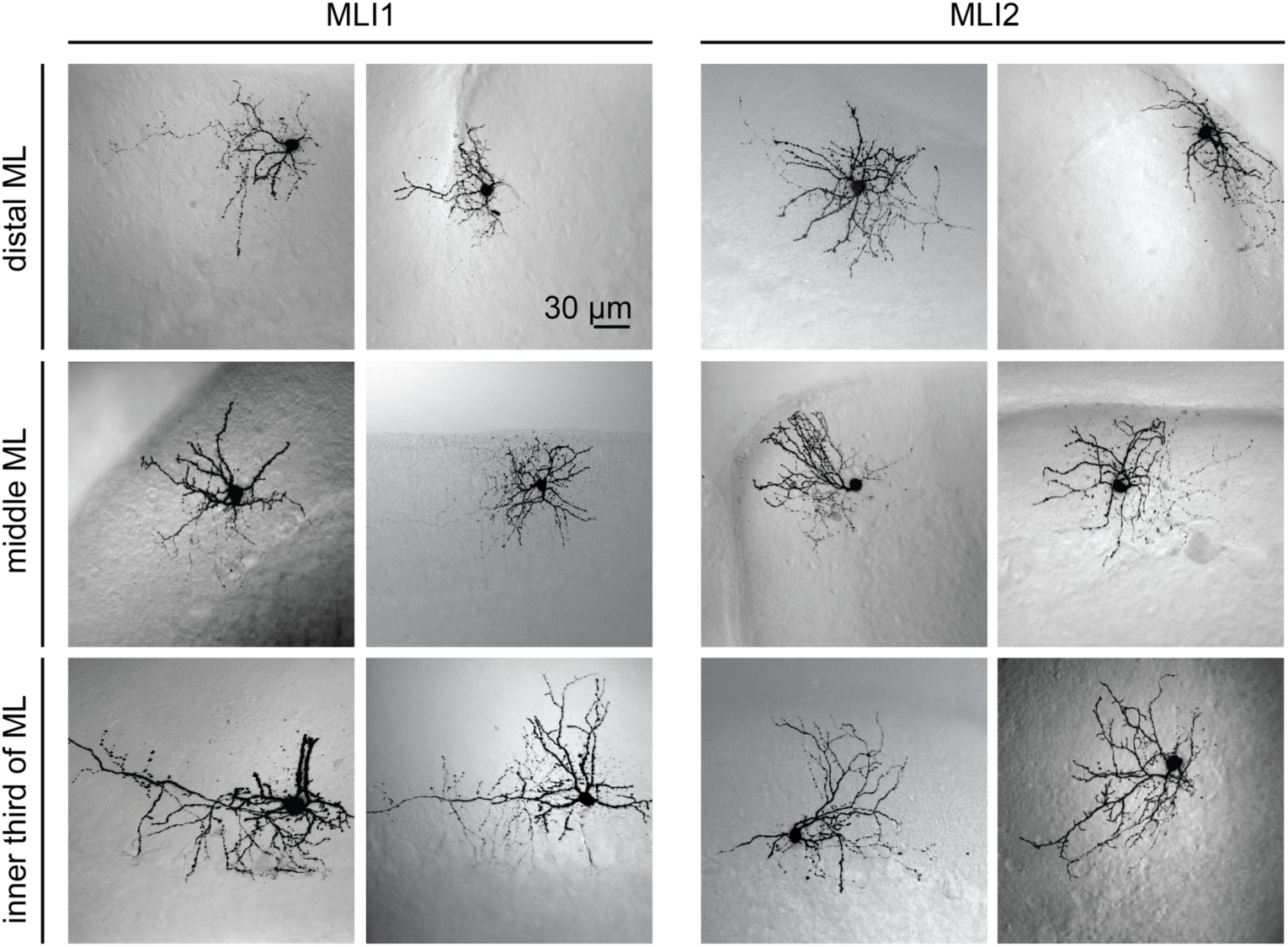
Additional 2-photon images of MLI1 and MLI2 neurons. MLIs were imaged and identified as in Fig. 4. Examples of MLI1s (left) and MLI2s (right) are shown for cells located in the distal third (top), the middle third (middle) and the inner third (bottom) of the molecular layer.

**Extended Data Figure 5:**
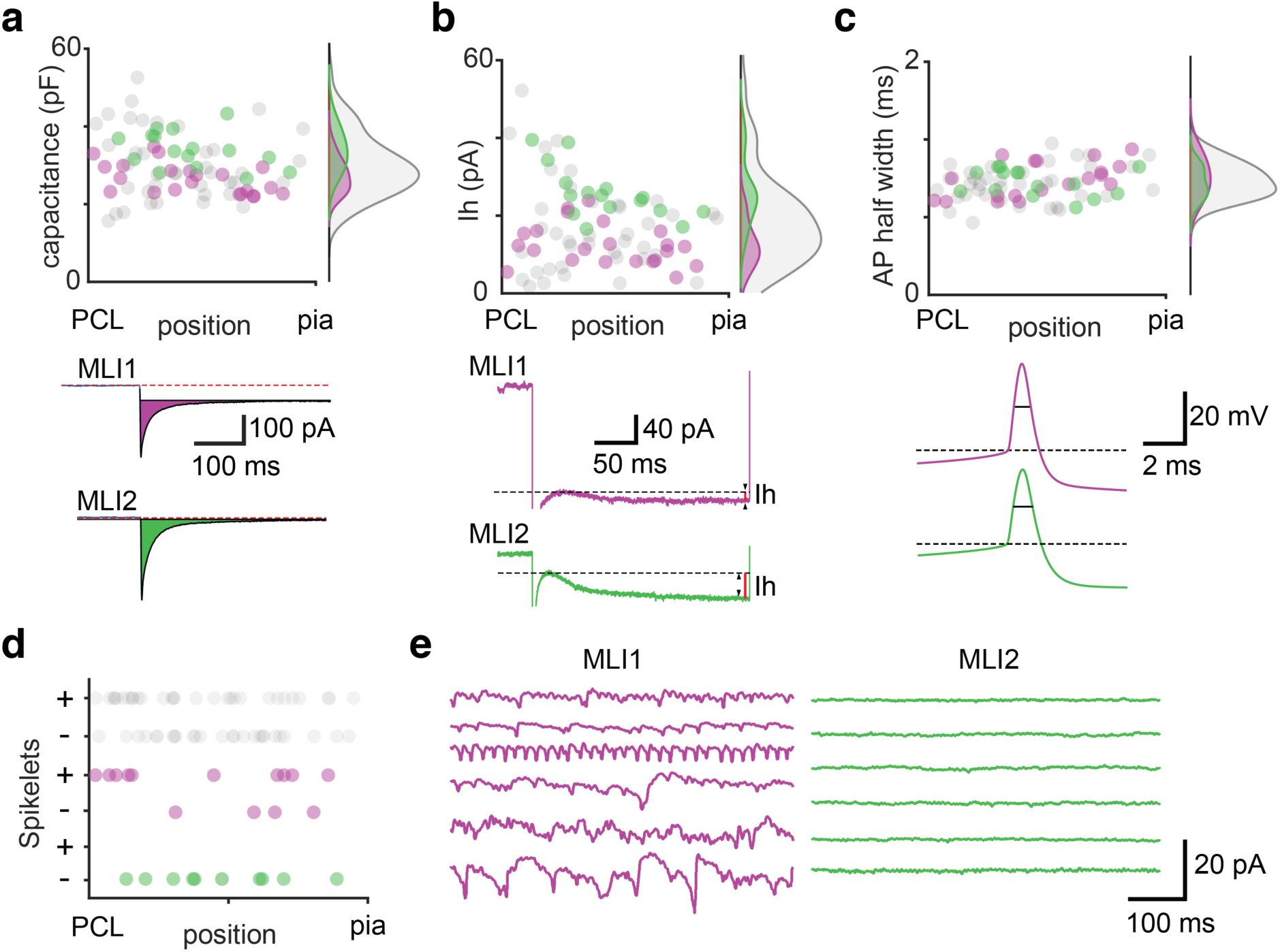
Additional comparison of electrical properties of MLI1 and MLI2 neurons. Measurements of: (**a**) Capacitance, (**b**) the amplitude of currents through hyperpolarization and nucleotide-gated (HCN) channels, I_h_, and (**c**) the action potential width in recordings of subsequently identified MLI1 (purple) and MLI2 (green) neurons. The properties are plotted as a function of position within the molecular layer. Density plots summarize the properties for all cells (grey), MLI1 (purple) and MLI2 (green). Capacitance was determined by measuring the responses to a -10 mV voltage step, and integrating the capacitive current as shown (purple and green shaded area for MLI1 and MLI2, respectively). These traces from cells in the inner third of the molecular layer show the large difference in resistance (R_m_= (−10 mV) I_SS_, where I_SS_ is the steady state current in response to the voltage step (black continuous line). Red dashed line denotes the baseline current. There was a significant difference in the capacitance of MLI1 and MLI2 neurons (p=2.03×10^−4^). (B, lower) The amplitude of I_h_ was determined by measuring the responses to a -30 mV step, and evaluating as shown. Measured I_h_ was significantly larger for MLI2 (p=4.24×10^−6^), and the difference was particularly striking for MLIs in the inner third of the molecular layer. (mC, lower) The action potential width was measured as shown and there was no significant difference for MLI1 and MLI2 neurons. **d**, The presence or absence of spikelets is shown as a function of position in the molecular layer is summarized for all MLIs, MLI1 and MLI2, with + indicating the presence of spikelets. **e**, Example recordings are shown for 6 MLI1 neurons (left, purple) and 6 MLI2 neurons (right, green). Number of cells and statistical tests are summarized in Extended Data Table 2.

**Extended Data Table 1:** Established cell type markers. Markers from existing studies used to assign clusters to specific established cerebellar cell type identities. Genes that lack a citation were used in annotation by examining the spatial localization in the AIBA^40^. Note that globular and candelabrum types could not be disentangled based upon the review of markers in the existing literature.

| <u>Cell Type</u> | <u>Defining marker(s)</u> |
| --- | --- |
| Purkinje | *Ppp1r17*<sup>34</sup> |
| Granular | *Gabra6*<sup>35</sup> |
| Golgi | *Slc6a5, Grm2, Sst*<sup>36</sup> |
| MLI1 | *Prkcd, Sorcs3, Ptprk* |
| MLI2 | *Prkcd, Nxph1, Cdh22* |
| PLI_3 (putative Lugaro) | *Htr2a*<sup>37</sup>, *Edil3* |
| PLI_1/PLI_2 (putative Globular/candelabrum) | *Aldh1a3, Slc6a5*<sup>37</sup> |
| Unipolar brush | *Eomes*<sup>38</sup> |
| Bergmann | *Gdf10*<sup>39</sup> |

**Extended Data Table 2:** Summary of electrophysiological data. Number of cells recorded for the data in Fig 4 and Extended Data Fig 5. Statistical tests and p-values are reported for each experiment. For analysis of input-output curves, a Poisson generalized linear mixed effect model of the form SpikeCount ∼ 1 + CurrentSteps + CellType + CurrentSteps*CellType + (1+CurrentSteps|CellIDs) with a canonical log link function was fit using maximum pseudo-likelihood estimation.

| Property | Figure | MLI1 n | MLI2 n | P-Value | Test | Total n |
| --- | --- | --- | --- | --- | --- | --- |
| Firing rate (spontaneous) | 4f (left) | 21 | 20 | 1.5E-03 | Mann Whitney test | 90 |
| R<sub>m</sub> | 4f (right) | 23 | 19 | 4.8E-06 | Mann Whitney test | 94 |
| IO curves (intercept) | 4g | 22 | 20 | 3.5E-05 | GLME |  |
| IO curves (slope) | 4g | 22 | 20 | 1.5E-04 | GLME |  |
| Spikelets | 4k | 10/14 | 0/10 | 5.8E-04 | Fisher's Exact test | 29/55 |
| Capacitance | E.D.5a | 23 | 19 | 2.0E-04 | Mann Whitney test | 94 |
| I<sub>h</sub> | E.D.5b | 23 | 19 | 4.2E-06 | Mann Whitney test | 93 |
| Spike width | E.D.5c | 20 | 19 | 3.1E-01 | Mann Whitney test | 78 |

